# MOZAIC: Compound Growth via *In Silico* Reactions and Global Optimization using Conformational Space Annealing

**DOI:** 10.64898/2026.03.07.710272

**Authors:** Jinhyeok Yoo, Woong-Hee Shin

## Abstract

**Motivation:** Fragment-based drug discovery (FBDD) is an efficient strategy that leverages small molecular fragments to explore broader chemical space by combining them. Advances in computational methods have enabled the calculation of molecular properties and docking scores, thereby accelerating the development of algorithm- and AI-based approaches in FBDD. However, it should be noted that certain methods do not provide synthetic pathways to obtain the proposed compounds. Consequently, these molecules might not be synthesized easily.

**Results:** In light of these developments, we propose MOZAIC, a novel framework that explores chemical space using a reaction-based fragment growing and Conformational Space Annealing, a powerful global optimization algorithm. Our results show that MOZAIC effectively produces chemically diverse molecules with balanced improvements in lead-like properties, including QED, synthetic accessibility, and binding affinity. Furthermore, its flexible objective function allows fine-tuning for specific design goals, such as enhancing solubility with binding affinity. These capabilities position MOZAIC as a valuable platform for advancing fragment-to-lead and lead optimization efforts in drug discovery.

**Availability and implementation:** MOZAIC is available at https://github.com/kucm-lsbi/MOZAIC/.

**Supplementary Information:** Supplementary data are available at *Bioinformatics* online.

## 1. Introduction

Fragment-based drug discovery (FBDD) has been successfully applied in drug discovery through exploring potent binding fragments to the subpocket of a receptor (Khedkar et al. 2025, Santos and da Silveira 2023, Xu and Kang 2025). These fragments generally yield higher hit rates than high-throughput screening due to their simple structures and drug-like properties (Hajduk and Greer. 2007). In addition, their diversity due to the combination allows cost- and time-efficient exploration of chemical space, even with relatively small libraries. The strategies of FBDD to optimize molecular properties and identify promising drug candidates can be categorized into three: growing, merging, and linking (Yoo et al. 2025). Fragment growing adds fragments from an initial one to improve molecular attributes, such as molecular size and solubility, with their selection guided by the binding pose in three-dimensional space and stability of growth. Fragment merging combines overlapping fragments that share a common substructure, producing a compound that preserves the core and incorporates both functional groups. Fragment linking connects fragments that bind to spatially distant subpockets or are chemically incompatible, using a molecular linker tailored to preserve their properties and support efficient binding.

Recent FBDD methods have evolved to address multi-objective tasks, achieving desirable properties simultaneously, such as drug-likeness and docking scores, through algorithmic and/or AI-driven approaches (Angelo et al. 2023). AutoGrow4 (Spiegel and Durrant 2020) implements a genetic algorithm (Forrest 1996) to generate diverse compounds from initial fragments. Subsequent to filtering generated molecules using Lipinski’s Rule of Five (Lipinski et al. 2012), the compounds are selected for the next generation, considering their docking scores and diversity. CReM-Dock (Minibaeva and Polishchuk 2024) utilizes fragments comprising C, N, O, S, and halogens derived from the ChEMBL database (Zdrazil et al. 2024). The fragments are filtered by applying BMS, Dundee, Glaxo, Inpharmatica, and PAINS filters, followed by synthetic accessibility (SA) filtration with a range of 2 to 2.5 (Ertl and Schuffenhauer 2009). The program generates molecules by connecting fragments, docking them using AutoDock Vina (Trott and Olson 2010) or GNINA (McNutt et al. 2025), and evaluating drug-likeness using the quantitative estimate of drug-likeness (QED) (Bickerton et al. 2012) within a genetic algorithm. FRAME (Powers et al. 2023) employs SE(3)-equivariant models to predict growth directions and fragments based on a reference protein-ligand complex. ReBADD-SE (Choi et al. 2023) is a reinforcement learning model using SELFIES (Krenn et al. 2022), incorporating a reward function that directs fragment growth based on target activity and drug-like properties. Despite the success of these programs in meeting the criteria of the FBDD, concerns have been raised regarding potential biases in the generated molecules. These concerns stem from the possibility that the genetic algorithm may be trapped around local minima, and from the potential for bias inherent in the training set when employing AI-based methods. Additionally, the synthesis of the proposed molecules might be challenging.

To address the challenges, we propose a novel fragment growing method, called MOZAIC (<u>m</u>olecule <u>o</u>ptimi<u>z</u>ation via <u>A</u>utoDock Vina, in silico fragment reaction, and <u>c</u>onformational space annealing). The fragments are added to the initial compound in accordance with the reaction rules defined by means of SMILES (simplified molecular input line entry system) Arbitrary Target Specification (SMARTS). By using the SMARTS rules based on the real organic reactions, the program would be able to provide synthesizable compounds, as well as the route to synthesize them. To produce molecules at the global optimum, Conformational Space Annealing (CSA) (Joung et al. 2018), a powerful population-based global optimization algorithm, was employed. CSA has been successfully applied to solve various biological problems, such as protein-ligand docking (Shin and Seok 2012, Shin et al. 2013), cryo-EM map fitting (Park et al. 2023), generation of molecules (Kwon and Lee 2021), and exploration of multiple reaction pathways (Lee et al. 2017). The key to CSA is the gradual narrowing of the distance cutoff when comparing the newly generated solutions and the current population, thereby providing diverse solutions. The objective function of MOZAIC is composed of three parts: The AutoDock Vina score is utilized to assess the binding affinity, while the SA score is employed to evaluate synthetic accessibility. Additionally, the QED score is employed to estimate drug-likeness. Furthermore, the components of the objective function could be readily substituted, enabling the generation of viable drug candidates that possess the desired molecular properties. This approach may offer a promising step toward bridging the gap between computational design and practical drug development using FBDD.

## 2. Materials and Methods

### 2.1 Overview of MOZAIC

MOZAIC is a fragment growing-based molecular optimization framework designed to generate target-specific, synthesizable compounds with drug-like properties. Figure 1 illustrates the flow of MOZAIC. The pipeline is composed of two phases. The initial ‘bank’ is constructed via SMARTS-based reactions (Figure 1A). In CSA, the term ‘bank’ denotes a set of ‘solutions’. Figure 1B illustrates the global optimization process utilizing CSA. In all phases, molecules are evaluated by an objective function, combining QED, SA, and Vina scores to balance synthesizability, drug-likeness, and binding potential to the target protein while exploring broad chemical space.

**Figure 1.**
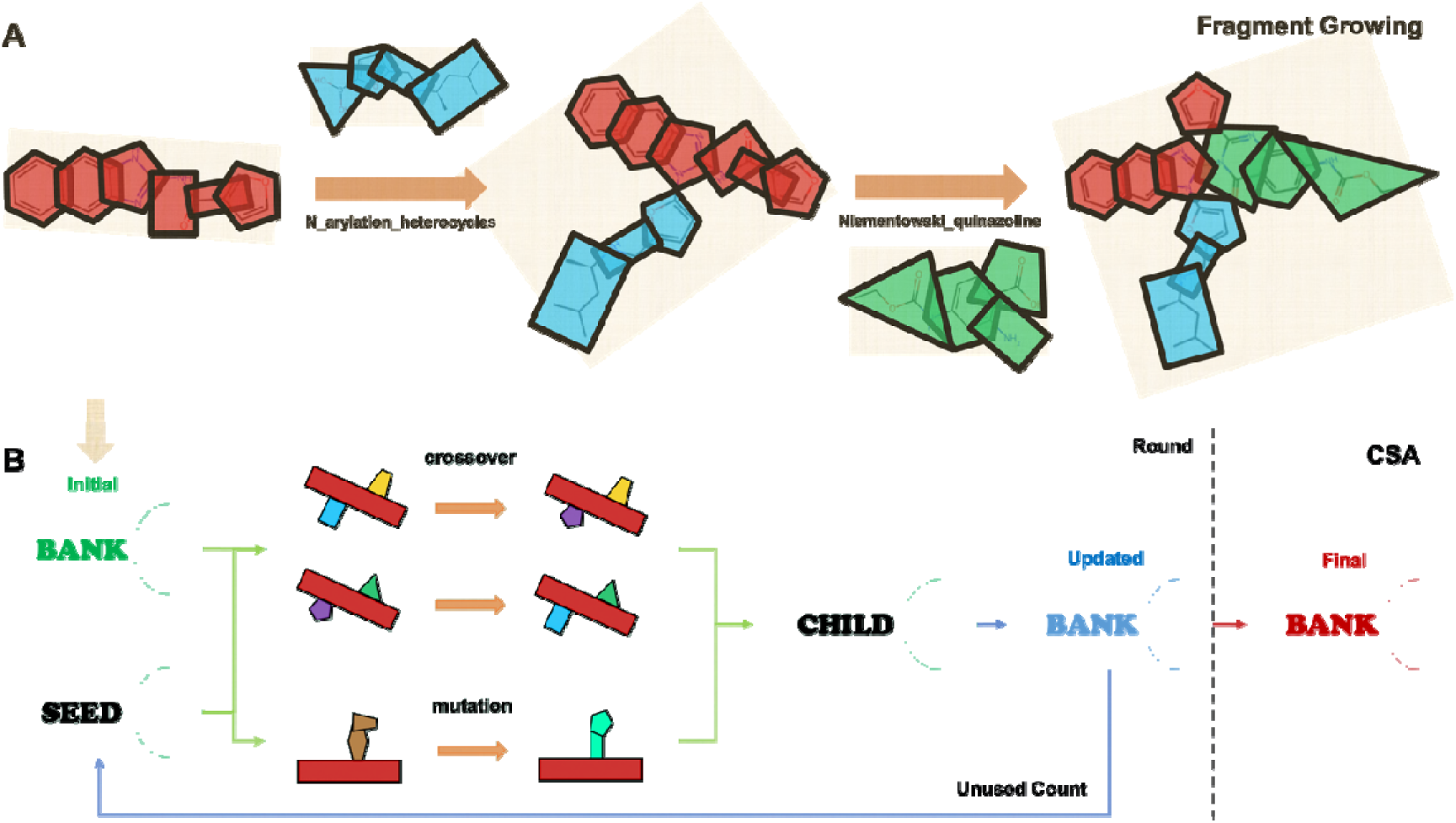
Overview of the MOZAIC framework. (**A**) Fragment growing by following in Silico reaction rules. The molecule is generated by adding fragments (blue and green) to the input compound (red) to construct initial molecules and children. (**B**) Optimization using CSA. From the current bank, seeds and partners are selected randomly to generate children via crossover and/or mutation. One round of CSA is ended if the number of unused seeds is less than a criterion. The final bank is obtained after three rounds of CSA.

The program’s input consists of a three-dimensional protein structure, residue indices that constitute a binding site, and a SMILES string representing a starting molecule. The construction of the initial bank entails the generation of new molecules (solutions) through the attachment of fragments to the starting compound, facilitated by predefined SMARTS-based reaction rules. Each solution record contains the following information: the functional group of the starting molecule, the reaction name and position (e.g., 65_Williamson_ether_0), and the SMILES strings of the added fragment and the resulting product. In the subsequent phase, ‘seed’ compounds are selected randomly from the current bank. The term ‘seed’ denotes that the compounds are used for operations to generate molecules. The new molecules, ‘children’, are generated by crossover or mutation, after selecting the partner molecules from the current bank. The children are compared to the current bank, considering molecular distances and objective function, followed by updating the current bank. This process is iterated three times, providing the final bank. The details of the CSA iteration will be given in the next subsection.

### 2.2 Application of CSA to molecule optimization by fragment-growing

#### 2.2.1 Objective function for evaluating the fitness of a molecule

The objective function in MOZAIC is composed of three metrics, QED, SA, and Vina score, to ensure the molecules have balanced properties. QED evaluates the drug-likeness of a molecule, based on eight physicochemical properties, ranging from zero to one. SA estimates synthetic accessibility on a scale of one to ten, with lower values indicating easy to synthesize. Vina score is a metric that predicts the binding affinity of a molecule to the target protein. A lower ΔG (kcal/mol) indicates stronger binding. To integrate these heterogeneous scores, the SA and Vina scores are scaled as Equations 1 and 2, respectively:

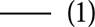

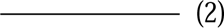

Following normalization, the three metrics are combined to form the objective function, which ranged from zero (worst) to three (best).

QED and SA scores are calculated using the RDKit (http://www.rdkit.org), while binding affinities and poses are obtained from AutoDock Vina (Trott and Olson 2010). After removing hetero-molecules, including water, from the input protein structure, a cubic docking box with a length of 20 Å is defined, centered at the geometrical center of the pre-selected binding site residues. The docking program generates ten poses, and the lowest score is used to calculate the objective function.

#### 2.2.2 Initial bank construction by applying in Silico reaction rules

The efficacy of CSA depends on the diversity and quality of the molecules comprising the initial bank (Lee et al. 2017). To get the diverse compounds, a candidate pool of 1000 molecules is constructed by adding fragments to the starting molecule using in Silico reaction rules, defined by SMARTS representation. A total of 104 reactions, composed of AutoClickChem (Durrant and McCammon 2012), Robust Rxn database (Hartenfeller et al. 2011), and ChemoDOTS set (Hoffer et al. 2024), are utilized to generate molecules. For the given input molecule, an examination is conducted of the atoms or bonds capable of forming reactions, as well as the types of reactions that could occur. Subsets of fragment databases are selected in accordance with the reaction rules. Subsequently, new molecules are generated by incorporating fragments selected randomly from the fragment library. To preserve the original scaffold, the maximum common substructure (MCS) between the initial and current compounds is computed. Atom indices of the initial functional groups are assigned to the MCS, then mapped onto the current compound to ensure that reactions occur only at the original reactive sites. The reaction history, containing the step, selected functional group, reaction position, and attached fragment, is recorded to perform crossover and mutation operations effectively during CSA. To promote molecular diversity even within the same scaffold, each reaction has a 10% chance of being skipped.

The MaxMinPicker algorithm is then employed to select 60 compounds from the candidate pool for the construction of the initial bank, with an emphasis on molecular diversity. The molecules are represented as 2048 bits by Extended Connectivity Fingerprints (ECFP) (Rogers and Hahn 2010) with a radius of four. Subsequently, the molecular distances are calculated as (1 – Tanimoto coefficient). After 60 compounds are selected for the initial bank, the half of the average of pairwise distance (*D_avg_*) was set to the initial value of *D_cut_*.

#### 2.2.3 New molecule generation, bank update, and CSA iteration

At each CSA iteration, a set of new molecules, referred to herein as children, is generated to explore a more extensive chemical space by employing crossover and mutation operations. Thirty seed molecules are randomly selected from the current bank. For the crossover procedure, two partner molecules are randomly selected, one from the initial bank and the other from the current bank, for each seed compound. The initial bank contributes to the preservation of diversity during the sampling process, while the current bank facilitates the exploration of emerging global minima of the objective function. If the paired seed and partner molecules have the same reaction functional group, the reaction information of the seed, including the added fragment, is replaced with that of the partner. The mutation re-selects the reaction components randomly based on the atom used for the reaction. In total, 90 compounds are generated and scored, 60 and 30 from crossover and mutation, respectively.

The generated child compounds are compared to the molecules in the current bank based on distance (*D*) and objective function. If the child is located within the *D_cut_* from any bank member, the closest pair is compared, and the current member is replaced if the child has a higher score. If the child compound has *D* larger than *D_cut_* from all current bank members, the compound with the lowest score is replaced by the child. The *D_cut_* is gradually reduced during CSA from *D_avg_*/2 to *D_avg_*/5. The use of *D_cut_* in CSA enables searching diverse space at the initial stage and focusing on the optimum solutions at the final steps. During each iteration, the selected seed molecules are marked as ‘used’, and in the next iteration, the seed compounds are selected from the ‘unused’ pool. A CSA round ends if the number of unused molecules is less than 30. The entire process of a single CSA round is repeated three times to construct the final bank.

#### 2.2.4 Final bank analysis and MOZAIC output

In addition to the features calculated in the objective function, the final bank includes the optimized molecules and their docking poses to the target receptor. The complexes are utilized for interaction analysis tools such as PLIP (Salentin et al. 2015), providing structural insights for further optimization. Thus, the final output comprises the information on both the initial and final bank compounds, including SMILES strings, scores, reaction details, and added fragments. Furthermore, target-ligand complexes are provided to facilitate 3D inspection of ligand binding conformations and subsequent interaction profiling.

### 2.3 Fragment library construction

Out of the 104 reactions, there are 89 bimolecular reactions. Consequently, a fragment library should be constructed in accordance with the established reaction rules. The construction process is illustrated in Supplementary Data Figure S1. From ZINC20 (Irwin et al. 2020) library (accessed July 2025), compounds that violate the Rule of Three (Jhoti et al. 2013) were filtered. The remaining compounds that matched the SMARTS patterns of the reaction rules were classified according to the reaction types. Considering the functional groups used in the reaction rules, 103 fragment subsets were constructed. When a reaction pattern matching produced more than 10,000 fragments, they were sorted by molecular weight, and 10,000 lowest-weight fragments were selected in ascending order. The total number of fragments is 893,675.

### 2.4 Benchmark set for MOZAIC

To evaluate the efficacy of the MOZAIC algorithm, three benchmark sets were prepared. The first benchmark was selected to assess the performance of MOZAIC. An examination of FBDD examples from the series of papers by Woodhead and his colleagues (Woodhead et al., 2024; Holvey et al., 2025) was conducted, leading the selection of two examples: Phosphodiesterase 10A (PDE10A) complexed with MK-8189 (PDB ID: 8DI4) and tRNA-(N^1^G37) methyltransferase (TrmD) co-crystalized with BDBM50613275 (PDB ID: 8APW). The selection was based on the number of functional groups in the initial molecule and the existence of a fully grown ligand complexed with the target protein in the PDB. The PDE10A was utilized to ascertain the impact of the quality of an initial bank on the CSA outputs. To provide a basis for comparison, the TrmD example was utilized in conjunction with CReM-Dock, a molecule generation method by fragment growing with a genetic algorithm and ligand docking. The AutoDock Vina parameters were kept the same as MOZAIC, and the chembl22_sa2_hac12.db fragment database was used. To further enhance sampling diversity, we used nclust = 2 for clustering and set max_replacements = 100.

In the second benchmark, MOZAIC was benchmarked on the protein structures that are not relevant to the starting molecule, to assess the robustness against receptor-structure. The top molecules from MOZAIC final bank were compared to the known active ligands. For this benchmark, beta-2 adrenergic receptor (ADRB2) was selected as a representative target. To select the starting molecule, compounds with pK_d_ or pK_i_ higher than seven were retrieved from the ChEMBL database (Gaulton et al. 2012). The filtered ligands were clustered using the Butina algorithm. For each cluster, the top five pK_i_ ligands were selected as cluster-specific references, and an MCS was derived from these ligands. To ensure consistency with the reaction space, we retained only clusters whose MCS contained at least two functional groups defined in the reaction SMARTS and selected the largest such cluster to maximize statistical support. The resulting MCS was used as the starting fragment for fragment growing. The overall process for starting fragment selection is illustrated in Supplementary Data Figure S2. For receptor structure, the carazolol-bound structure (PDB ID: 2RH1, resolution: 2.40 Å), which was used in structure-based docking and virtual screening studies (Kolb et al. 2009). The Tanimoto similarity between carazolol and the starting compound is 0.091. We also tested MOZAIC against the AlphaFold3 (AF3) structure (Abramson et al. 2024). The predicted structure was superposed onto the 2RH1 template using UCSF Chimera MatchMaker (Meng et al. 2006), resulting in an RMSD of 0.746 Å over 439 pruned atom pairs. Based on this alignment, binding site residues in the AF3 model were defined as those located within 5 Å of the carazolol ligand in the crystal structure. The local confidence of these residues was quantified by atom-averaged pLDDT scores, yielding a mean binding site pLDDT of 87.41, with 8 out of 20 residues showing a pLDDT more than 90.

The part of MOZAIC objective function, such as QED, can be easily substituted with alternative molecular descriptors, thereby providing flexibility in the optimization of multi-objectives in molecular design. To demonstrate this, the program was utilized to improve the solubility of Peroxisome Proliferator-Activated Receptor gamma (PPARγ) binding ligands. The protein belongs to the nuclear receptor family, and acts as a key metabolic switch for regulation of adipocyte differentiation and glucose homeostasis (Houseknecht et al. 2002). For nuclear receptors, many potent ligands are relatively lipophilic (Zhang et al. 2015); therefore, improving aqueous solubility while maintaining or even improving affinity is critical for oral exposure, formulation flexibility, and predictable pharmacokinetics. To enhance solubility, MOZAIC objective function was modified as a combination of molecular solubility and docking score. The solubility of a molecule is predicted by ESOL (Delaney 2004), a linear regression model (Equation 3).

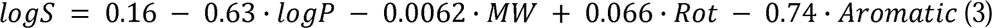

where *MW*, *Rot,* and *Aromatic* represent molecular weight, the number of rotatable bonds, and the proportion of aromatic atoms, respectively. The ESOL scores were normalized same as the AutoDock Vina score.

From ChEMBL, the lowest molecular weight compound with pChEMBL higher than eight was selected as the initial molecule to be optimized: CHEMBL375270 (EC_50_: 2.0 nM). The PDB ID 6MS7 with a resolution of 1.43 Å was selected for the PPARγ structure.

## 3. Result and discussion

### 3.1 Applying MOZAIC to optimize PDE10A hit compound and comparing initial bank construction methods

To validate the performance of MOZAIC, PDE10A was selected as a first example. 4,6-dichloro-2-cyclopropyl-5-methylpyrimidine with 8.7 μM of K_i_ to the target protein was employed as the initial molecule for the benchmark. The molecule was fully optimized to MK-8189 (Layton et al. 2023), a selective PDE10A inhibitor with K_i_ = 0.029 nM. MOZAIC was performed on the co-crystalized structure of the receptor and MK-8189.

As illustrated in Figure 2, the program demonstrates notable efficacy in terms of general optimization capability. Figure 2A compares the two-dimensional structures and property values of the initial molecule, MK-8189, and the best compound from the final bank. The best compound was generated via two C–C coupling steps: a Heck reaction between a vinyl halide and an alkene, followed by a Negishi coupling. The predicted binding affinity of the molecule exhibited a substantial enhancement from the initial compound. It was improved by about 70%, shifting from -6.038 kcal/mol to -10.359 kcal/mol. Notably, this value is also higher than the lead compound, MK-8189 (-7.770 kcal/mol). Although QED and SA either remained or deteriorated due to the increased complexity by adding fragments, these values remain comparable to those of MK-8189. Figure 2B illustrates the binding poses and PLIP-derived interaction profiles of the three complexes. Interestingly, the MOZAIC best compound and MK-8189 share similar binding interaction patterns with the receptor. PLIP identified that V668, I682, Y683, F686, P702, E711, Q716, and F719 are common interacting residues in both binding poses. The pyrimidine core of both molecules forms a π-stacking interaction with F719 and a hydrogen bond with Q716. The selectivity of PDE10A comes from the ‘selectivity pocket’ (Bhardwaj and Purohit 2021) which is not conserved in other PDE family proteins. The 5-methyl-pyridine of MK-8189 has been observed to penetrate the selectivity pocket of the protein, where it has been shown to interact with Y683 through the formation of a hydrogen bond. The MOZAIC optimized compound also forms an interaction between the terminal pyridine and the tyrosine by hydrophobic interaction. Compared with the initial pose, MK-8189 and the final compound occupy the binding pocket more broadly, engaging a greater number of nearby residues. These observations indicate that the binding coverage and interaction density have been enhanced, which may contribute to improved molecular recognition and stability (Bissantz et al. 2010, Myslinski et al. 2011).

**Figure 2.**
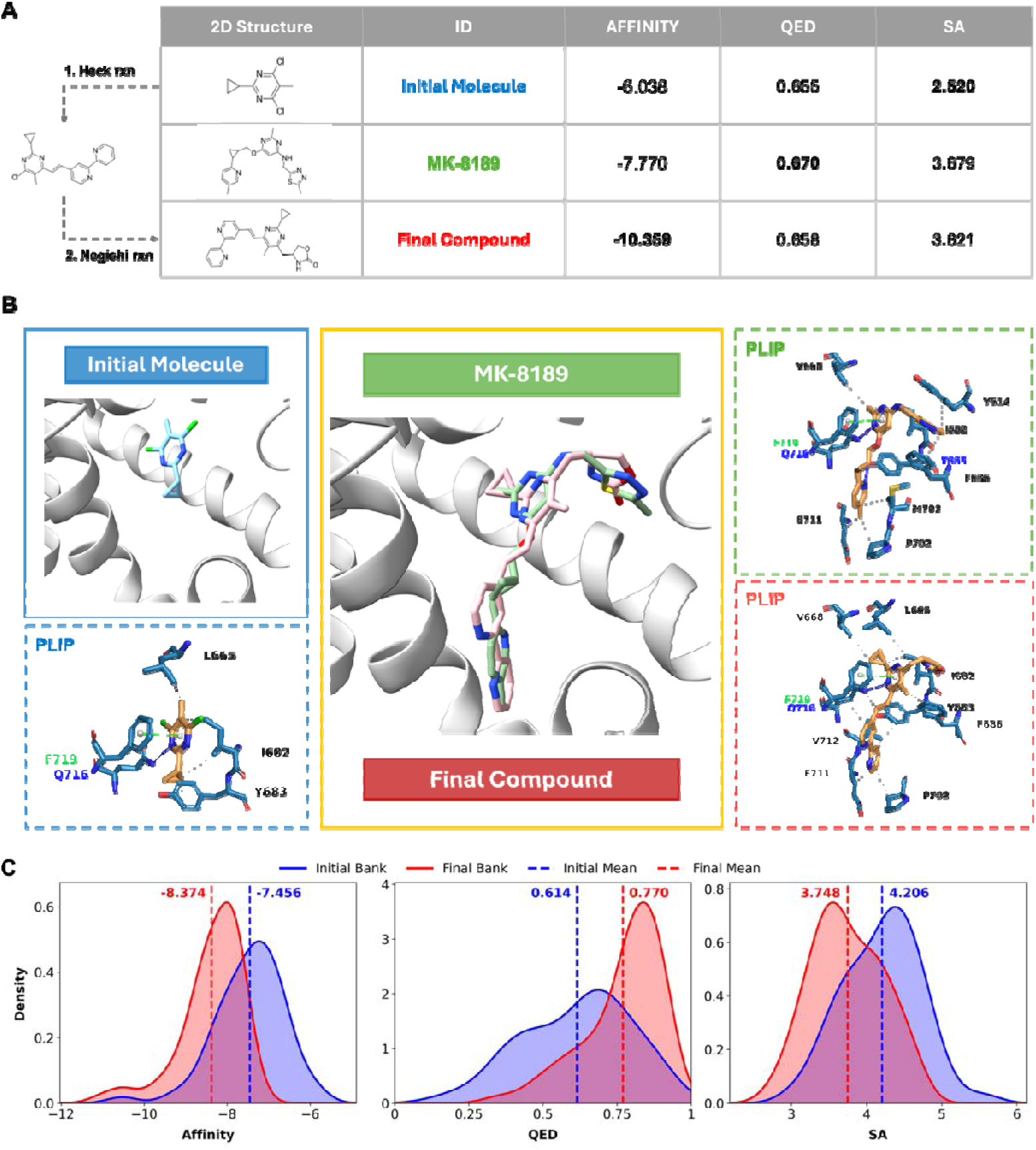
(**A**) Comparison of the initial molecule, MK-8189, and the best compound from the final bank in terms of 2D structures and predicted properties. (**B**) Binding poses and PLIP-derived interaction profiles of the three complexes. (**C**) Distributions of predicted binding affinity (left), QED (middle), and SA scores (right) for initial and final banks. The blue and red solid lines show the property dispersals of the initial bank and final bank, respectively. The dashed lines represent the average values of the properties.

The distributions of three molecular properties of the initial bank and the final bank are presented in Figure 2C: AutoDock Vina score, QED, and SA score. All metrics shifted toward improved values. The mean values of binding affinity, QED, and SA shifted from - 7.456 kcal/mol, 0.614, and 4.206 to -8.374 kcal/mol, 0.770, and 3.748, respectively. It demonstrates MOZAIC’s capacity to optimize multiple objectives in a balanced manner simultaneously.

As CSA is one of the population-based global optimization algorithms, its performance depends on the initial population (Shin et al. 2013). To investigate how the initial bank affects the performance of MOZAIC in the CSA process, a comparison of three different selection strategies for constructing the initial bank was conducted: MaxMinPicker of RDKit considering molecular diversity, score-based selection method prioritizing top-ranked compounds (QED + SA), and random sampling. Initially, a pool of 10,000 molecules was generated, and 1,000 compounds were selected through the application of three distinct methods. Figure 3 shows the differences between the selection methods.

**Figure 3.**
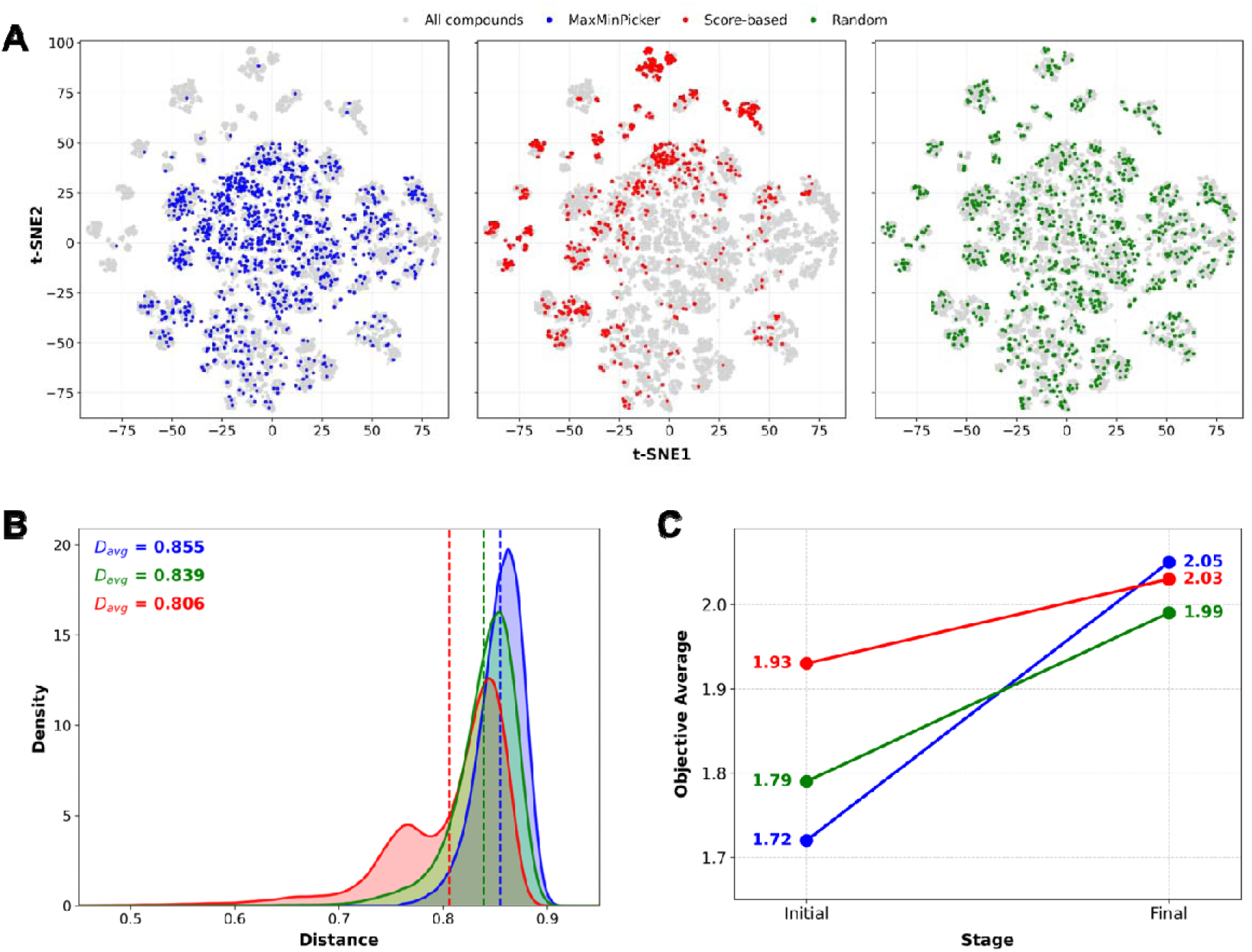
(**A**) A t-SNE projection of initial BANKs selected by three methods: MaxMinPicker (left, blue), Top-score selection (middle, red), and random selection (right, green). The generated 10,000 compounds are colored grey. (**B**) Pairwise distance distributions (solid line) and average distance (dashed line) of each initial bank compound. (**C**) Change in the average scores of the initial and final banks for each strategy. The MaxMinPicker, top-score selection, and random selection are colored blue, red, and green, respectively.

Figure 3A illustrates the two-dimensional projection of pool compounds’ chemical space. The ECFP fingerprint with a radius of four and 2048 bits was calculated for the pool compounds and further embedded in two-dimensional space using t-SNE. As anticipated, MaxMinPicker identified the molecules exhibiting the highest degree of diversity, with random selection and score-based selection following in succession (Figure 3B). The mean average distance of the initial bank is observed to be the highest for the MaxMinPicker (0.855), suggesting that it attained the most heterogeneous set in the original chemical space. Conversely, the score-based selection method identified compounds within a more constrained chemical space, exhibiting an average value of 0.806. The random selection is situated in the median position, with a mean value of 0.839. To evaluate the downstream impact, MOZAIC was executed with the three different initial banks, and the average objective scores of the initial and final banks for each selection method were calculated (Figure 3C). In this experiment, an initial pool of 1,000 molecules was generated, and 60 of them were selected to form the initial bank, using the three selection methods. While score-based selection started with the highest initial average, its improvement was modest. In contrast, MaxMinPicker, despite starting with the lowest initial average, led to the largest increase in objective score and yielded the highest final average. These results demonstrate that a diverse initial bank might promote better global optimization performance.

*3.2 Optimizing a compound targeting TrmD and comparing to CReM-dock*

In order to assess the performance of MOZAIC and to compare it with other molecule optimization methods, TrmD was selected as the next benchmark. CReM-dock approach was selected for comparison because it utilizes AutoDock Vina and a genetic algorithm to add the fragments, similar to MOZAIC. Indole-5-carboxamide, with an IC_50_ of 160 μM to the target protein, was selected as the initial molecule. Wilkinson and colleagues (Wilkinson et al. 2023) optimized the molecule to BDBM50613282, with an IC_50_ of 1.3 nM.

Table 1 shows the top three molecules and their properties obtained by the two programs, along with the initial molecule and BDBM50613282. All three molecules from MOZAIC exhibit stronger predicted binding affinities with a mean of -9.683 kcal/mol compared to the initial molecule (-6.891 kcal/mol), and even lower than the optimized molecule (-8.220 kcal/mol). Conversely, the SA score and QED demonstrate a similarity to, or in some cases, a lower performance than that of the optimized compound. CReM-dock also successfully produced the molecules exhibiting lower binding affinities compared to the initial compound and the optimized one, with an average binding energy of -8.428 kcal/mol.

**Table 1.**
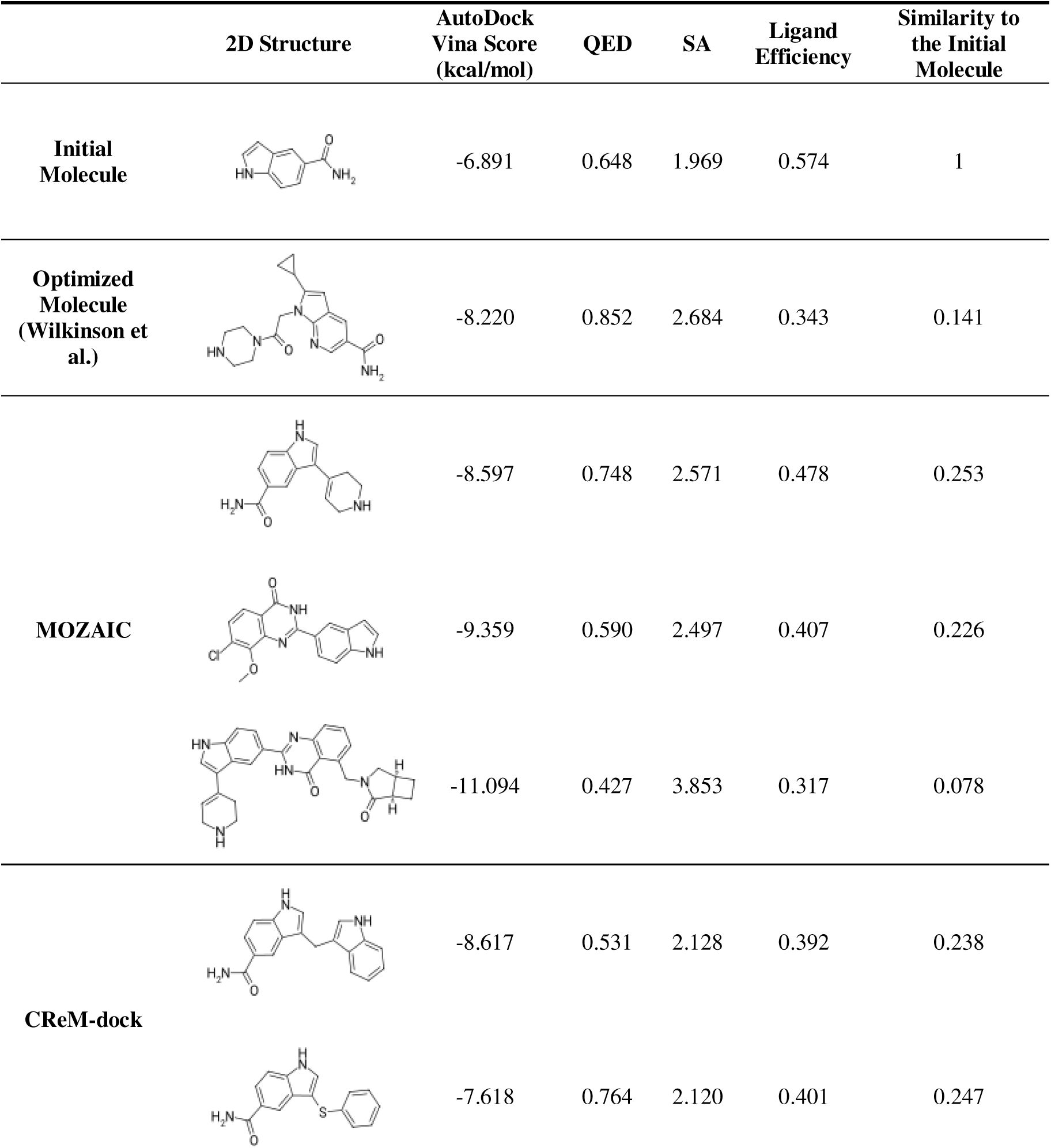
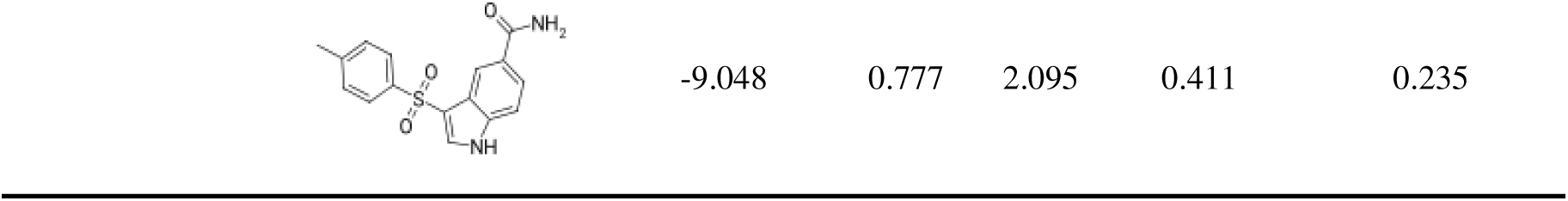
The top three compounds from CReM-dock and MOZAIC, compared by two-dimensional structures and properties.

|  | 2D Structure | AutoDock<br>Vina Score<br>(kcal/mol) | QED | SA | Ligand<br>Efficiency | Similarity to<br>the Initial<br>Molecule |
| --- | --- | --- | --- | --- | --- | --- |
| **Initial**<br>**Molecule** |  | -6.891 | 0.648 | 1.969 | 0.574 | 1 |
| **Optimized**<br>**Molecule**<br>**(Wilkinson et**<br>**al.)** |  | -8.220 | 0.852 | 2.684 | 0.343 | 0.141 |
| **MOZAIC** |  | -8.597 | 0.748 | 2.571 | 0.478 | 0.253 |
|  |  | -9.359 | 0.590 | 2.497 | 0.407 | 0.226 |
|  |  | -11.094 | 0.427 | 3.853 | 0.317 | 0.078 |
| **CReM-dock** |  | -8.617 | 0.531 | 2.128 | 0.392 | 0.238 |
|  |  | -7.618 | 0.764 | 2.120 | 0.401 | 0.247 |

A comparison of the top three molecules from each method reveals that MOZAIC produced stronger and larger binders than CReM-dock. The docking score is predicated on an atom-pairwise additive method that predicts binding affinity, thus resulting in a large molecule having low binding affinity. Therefore, the ligand efficiency was calculated by dividing the absolute value of the predicted score by the number of heavy atoms to reduce the effect of the molecule size. The arithmetic mean of the top three molecules from each program yielded the same value: 0.401. On the other hand, CReM-dock showed better values for QED and SA than MOZAIC. The average values of CReM-dock are 0.691 and 2.114, while those of MOZAIC are 0.588 and 2.974 for QED and SA, respectively. However, the top three CReM-dock molecules show minimal variation—primarily small substituents added to the same positions—resulting in limited molecular diversity. In contrast, MOZAIC generates structurally diverse compounds, with examples that include ring formation and distinct modifications. Consistent with this observation, the final 60 compounds of MOZAIC exhibit greater divergence from the initial molecule. The average Tanimoto similarity to the initial molecule is 0.135, ranges from 0.074 to 0.253, whereas the top 60 CReM-dock compounds remain closer to the initial structure, from 0.198 to 0.412, with an average of 0.285. This increased deviation is accompanied by broader chemotype coverage. Based on Bemis-Murcko scaffolds, the MOZAIC approach generates a larger number of unique scaffolds with higher scaffold entropy. The number of unique scaffolds generated by MOZAIC and CReM-dock are 45 and 32, and the entropies are 3.591 and 2.791, respectively. Also MOZAIC shows a lower dominance of the most frequent scaffold, 0.133, while CReM-dock exhibits 0.367. These findings suggest that MOZAIC searches a broader exploration of chemical space. In addition, while CReM-dock expands fragments by attaching new moieties to available hydrogen atoms, MOZAIC performs reaction-based modifications guided by SMARTS rules. This allows MOZAIC to mimic more realistic synthetic transformations, could yield products with improved synthetic feasibility and broader chemical novelty.

### 3.3 Optimizing compounds in the absence of relevant receptor structures

Since MOZAIC employs AutoDock Vina, its performance might be sensitive to the receptor conformation. To investigate the dependency, the program was started using the co-crystal structure with the ligand not similar to the starting molecule and AF3-predicted structures. Table 2 shows the top molecules optimized by MOZAIC, the starting fragment, and the top five pK_i_ compounds from the same cluster of the starting molecule.

**Table 2.**
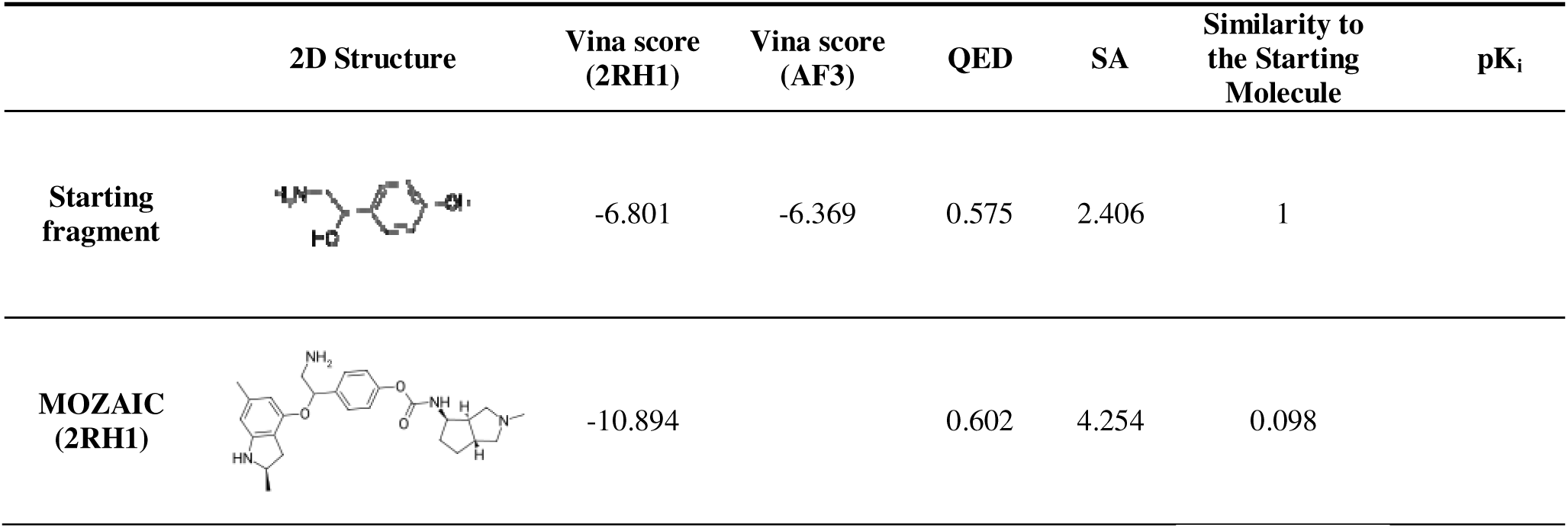
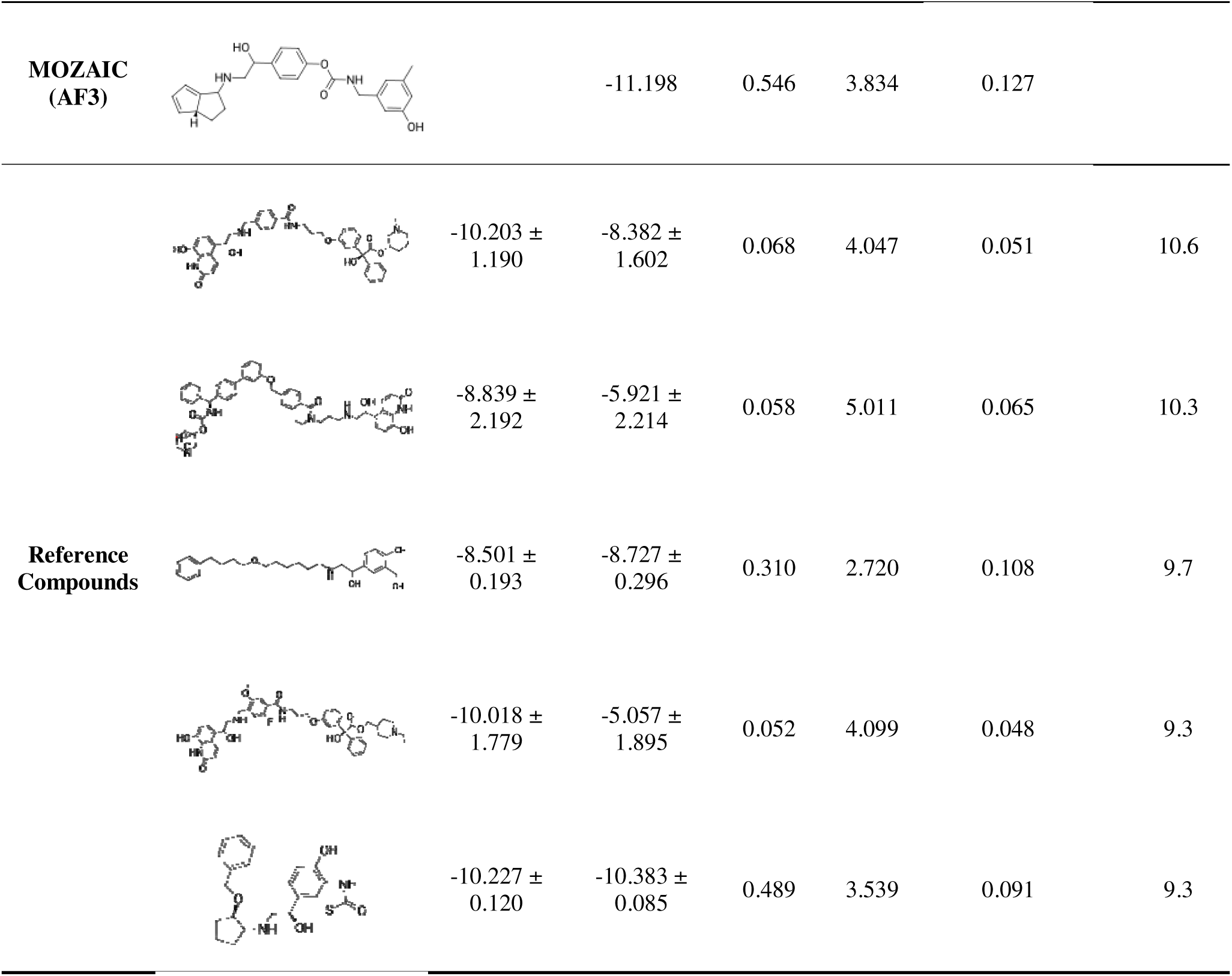
Comparison of the final compounds optimized by MOZAIC with predefined reference ligands on the ADRB2 crystal structure (PDB ID: 2RH1) and AF3 structure.

The best MOZAIC optimized compound was obtained through Mitsunobu reaction to convert alcohol at the phenol site to ester and isocyanate reaction with alcohol. The compound showed competitive values to the reference compounds, with pK_i_ values higher than nine, in terms of Vina scores. On the other hand, the MOZAIC compounds exhibited more favorable QED profiles and comparable SA scores than the references. In addition, the best MOZAIC compounds showed a low similarity to the starting compound, comparable to the references, indicating that they have evolved into structurally distinct entities.

Initially, MOZAIC identified three reaction sites from the starting molecule: hydroxyl, primary amine, and phenol groups. Most of the initial bank compounds for the X-ray structure substituted all functional groups to fragments. Out of 60, 45 molecules were tri-site reacted ones for the crystal structure and AF3 structure, respectively. However, the majority of the final banks were composed of bi-site reacted molecules, 43 out of 60 compounds for both structures to balance the binding affinity and other two features. On average, the molecules have binding affinity of -9.391 kcal/mol, while 0.672 and 3.692 for QED and SA score. The three uni-site reacted compounds in the final bank yield favorable QED (average: 0.798) and SA scores (average: 3.247), but limited affinity with an average of -8.437 kcal/mol. On the other hand, tri-site substitution can enhance affinity while losing drug-likeness and synthesizability, -9.708 kcal/mol, 0.650, and 4.188 for AutoDock Vina score, QED, and SA, respectively.

Similar to the X-ray structure, the AF3 structure-based optimized molecule showed a comparable binding affinity and improved QED and SA scores to the reference molecules. The reference molecules showed relatively weaker binding affinities when docked to the AF3-predicted structure. This suggests MOZAIC is capable of identifying diverse chemotypes that retain docking performance even under receptor-structure uncertainty.

### 3.4 Customization of Physicochemical Objectives in MOZAIC

Since the components of MOZAIC objective function are coded in Python3 and modularized, they can be easily replaced. Thus, the program can optimize physico-chemical features of compounds other than AutoDock Vina, QED, and SA score. We replaced QED and SA score with ESOL, a simple regression model that predicts logS of compounds, and selected a ligand from the ChEMBL database that binds to PPARγ, to assess whether MOZAIC can improve both solubility and binding affinity simultaneously. The starting compound selected from the ChEMBL database and the top three molecules generated by MOZAIC are shown in Table 3.

**Table 3.**
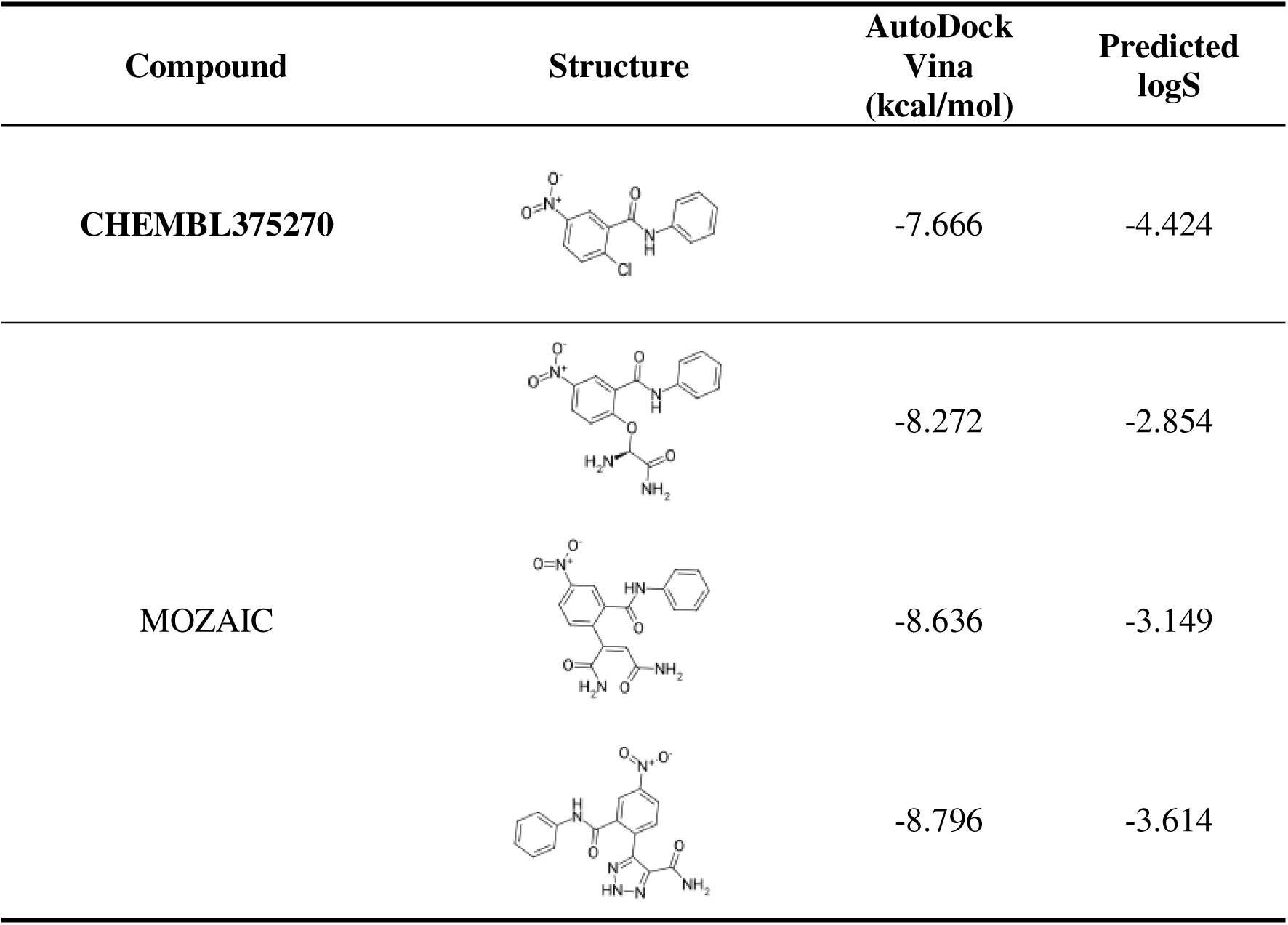
The starting compound selected from ChEMBL that binds to PPARγ and its top three optimized compounds by MOZAIC.

| Compound | Structure | AutoDock Vina<br>(kcal/mol) | Predicted<br>logS |
| --- | --- | --- | --- |
| CHEMBL375270 |  | -7.666 | -4.424 |
| MOZAIC |  | -8.272 | -2.854 |
|  |  | -8.636 | -3.149 |
|  |  | -8.796 | -3.614 |

MOZAIC identified two potential reaction sites of given starting molecule, amine/amide and halide. Except for one compound with uni-site reaction, the initial bank was composed of bi-site reacted compounds. However, the trend is opposite in the final bank. The only two molecules out of 60 were bi-site reacted compounds, while the others are uni-site substituted ones. This shift appears to be driven by the specific topology of the Y- or T-shaped binding pocket in PPARγ (Kroker and Bruning 2015), where dual-site expansion likely caused steric clashes and excessive hydrophobicity. Indeed, the initial bank exhibited poor solubility, with a mean logS of -8.563 (±1.278) and high Vina score variance (-7.797 ± 2.628), whereas the final solutions achieved a stable high-Vina score profile (-8.599 ± 0.561, range: -10.540 to -7.612) with significantly improved solubility (-4.647 ± 0.593).

The top three-optimized molecules replaced the chlorine atom attached to the para-position of the nitro group to a fragment using Halide_and_Alcohol or Stille reactions. Comparison with the starting molecule revealed that both AutoDock Vina score and logS were enhanced. The mean values of the three compounds are -8.568 kcal/mol and -3.206 for AutoDock Vina score and logS, respectively. The modifications increased molecular polarity while improving the predicted binding affinity (Supplementary Data Figure S3). Notably, the MOZAIC compounds maintain the nitro position observed in the initial molecule, while redirecting the para substituent into the interior subpocket. This reorientation increases volumetric occupancy and shape complementarity within the confined Y-shaped cavity, leading to tighter packing in the buried region and a more consistent pose family across the optimized derivatives. Consequently, the program successfully optimized the initial molecule to identify solution candidates, achieving a favorable balance between enhanced solubility and binding potency.

## 4. Conclusion

In this study, we presented MOZAIC, a structure-based fragment growing framework that integrates reaction rules with the CSA algorithm. A powerful global optimization algorithm, CSA, enables MOZAIC to rigorously explore the chemical space defined by synthetic rules, ensuring both optimization quality and structural diversity. The application of this framework to benchmarks demonstrated its advantages. First, MOZAIC could suggest potent compounds comparable to the lead compound in terms of the docking score. Second, facilitated by CSA, MOZAIC produced compounds with higher scaffold entropy and lower structural redundancy, effectively avoiding the generation of repetitive analogues. Third, the program identified alternative chemotypes that retained favorable docking scores, demonstrating robustness against receptor-structure uncertainty. Finally, it showed a flexibility in choosing physico-chemical properties to be optimized.

However, there are several aspects to be improved. The current implementation relies on AutoDock Vina, which has limited accuracy in absolute quantitative affinities, as it does not adequately account for target flexibility or solvent effects (Pantsar and Poso 2018).

Additionally, the chemical space is explored via the predefined reaction rules, restricting molecular diversity (Saldívar-González et al. 2020). To address these limitations, the future work will focus on expanding the accessible chemical space by integrating a broader set of SMARTS-based reactions. Advanced scoring methods, such as quantum mechanics-based potentials or Vina-GPU (Tang et al. 2024), will be incorporated into the scoring function to enhance accuracy and speed. For blind docking scenarios, tools such as Fpocket (Le Guilloux et al. 2009) and P2Rank (Krivák and Hoksza 2018) could be incorporated for binding site prediction.

## Acknowledgement Funding Information

This work was supported by the National Research Foundation (NRF) funded by the Korea government (Ministry of Science and ICT) [RS-2024-00441029].

## Data Availability

All benchmarked data is available at https://github.com/kucm-lsbi/MOZAIC/tree/main/benchmark_results/.

## Code Availability

The source code for MOZAIC is available at https://github.com/kucm-lsbi/MOZAIC/.

