## Supplementary Data for "MOZAIC: Compound Growth via *In Silico* Reactions and Global Optimization using Conformational Space Annealing"

**Fragment Library Construction Process**

1. We downloaded ‘Annotated’ molecules from ZINC20 and applied initial filters of logP < 3 and molecular weight < 300, resulting in ~450 milion SMILES entries.

2. Physicochemical properties of entries were calculated using RDKit, including the number of hydrogen bond donors/acceptors and rotatable bonds, and further filtered according to the Rule of Three (Ro3) (Jhoti et al. 2013) for fragments.

3. Redundant entries were removed using canonical SMILES, duplicates were collapsed into single representatives, leaving approximately 80 million unique fragments.

4. Finally, fragments containing functional groups required for AutoClickChem, Robust and ChemoDOTS reactions were selected, and the final library was constructed with up to 10,000 fragments, ordered by increasing molecular weight.

**Figure S1.** Process of constructing the fragment library

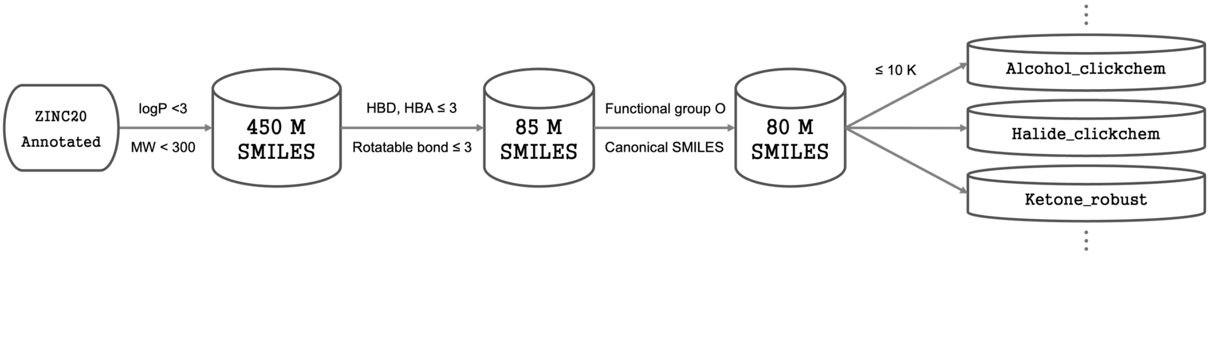

**Selecting the starting fragment from ChEMBL for ADRB2**

1.Collect active compounds with pChEMBL ≥ 7; 391 compounds are retained (for duplicate SMILES, only the entry with the highest pChEMBL is kept).

2. Calculate ECFP fingerprint with radius of 4, 2048 bits.

3. Clustering ligands using Butina algorithm with a Tanimoto coefficient of 0.7.

4. Derive an MCS from the top-5 pChEMBL ligands in each cluster, and use the MCS from the largest selected cluster as the starting fragment.

**Figure S2.** Workflow for starting fragment extraction.

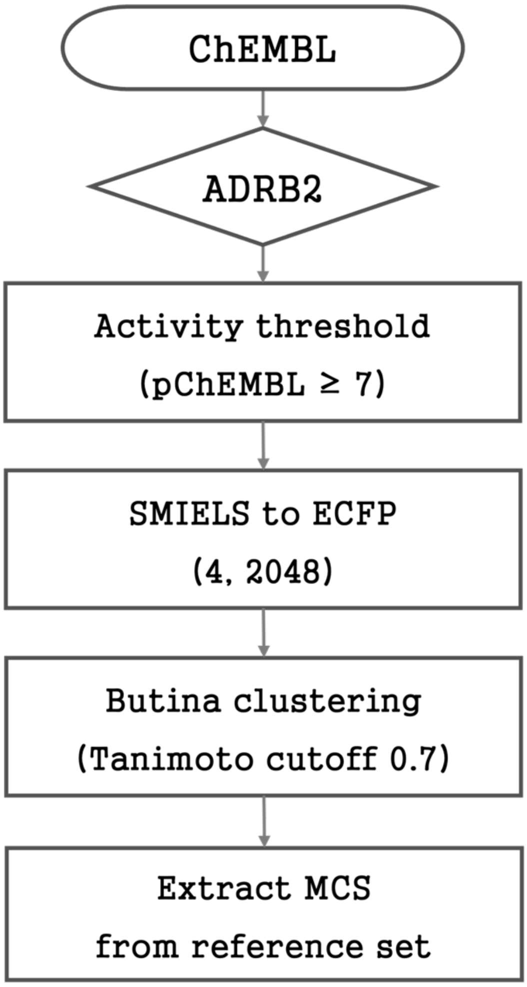

**Figure S3.** Superposed docking poses of the initial compound (sky blue) and MOZAIC derivatives (pink shades), illustrating pose convergence among optimized compounds.

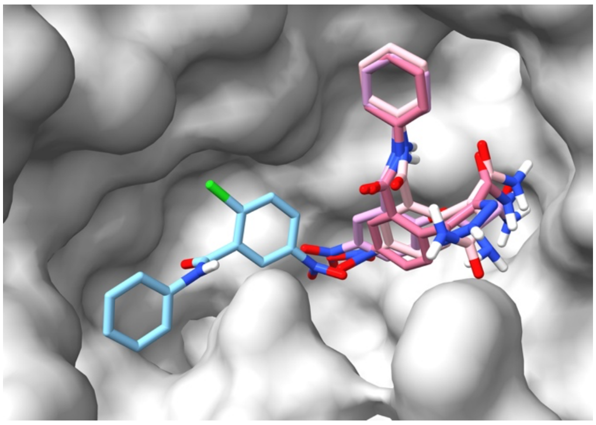
